# SSNA1 stabilizes dynamic microtubules and detects microtubule damage

**DOI:** 10.1101/2021.02.05.429994

**Authors:** EJ Lawrence, C Arnaiz, G Arpag, M Zanic

## Abstract

Sjögren’s Syndrome Nuclear Autoantigen 1 (SSNA1/NA14) is a microtubule-associated protein with important functions in cilia, dividing cells and developing neurons. However, the direct effects of SSNA1 on microtubules are not known. We employed *in vitro* reconstitution with purified proteins and TIRF microscopy to investigate the activity of human SSNA1 on dynamic microtubule ends and lattices. We find that SSNA1 modulates all parameters of microtubule dynamic instability – slowing down the rates of growth, shrinkage and catastrophe, and promoting rescue. SSNA1 accumulation on dynamic microtubule ends correlates with the growth rate slow-down. Furthermore, SSNA1 prevents catastrophe when soluble tubulin is removed or sequestered by Op18/Stathmin. Finally, SSNA1 detects spastin-induced damage and inhibits spastin’s severing activity. Therefore, SSNA1 is both a potent microtubule stabilizing protein and a sensor of microtubule damage; activities that likely underlie SSNA1’s cellular functions.

## Introduction

Sjogren’s syndrome nuclear autoantigen-1 (SSNA1/NA14) is a microtubule-associated protein (MAP) that plays important roles in cilia, cell division and neuronal development. In cilia, SSNA1 localizes to basal bodies and axonemes where it is required for proper cilium assembly and intraflagellar transport (Pfannenschmid et al. 2003, Schoppmeier et al. 2005, Lai et al. 2011). In dividing cells, SSNA1 is enriched at the spindle poles and midbody, and is necessary for proper cell division (Pfannenschmid et al. 2003, Goyal et al. 2014). Finally, SSNA1 promotes axon elongation and branching in developing neurons (Goyal et al. 2014, Basnet et al. 2018). Although SSNA1 is involved in a range of microtubule-driven cellular processes, its direct effects on microtubules are not known.

SSNA1 is a small (∼14 kDa), coiled-coil protein that self-assembles into higher-order filaments (Ramos-Morales et al. 1998, Rodriguez-Rodriguez et al. 2011, Basnet et al. 2018). A recent *in vitro* study using cryo-EM/ET reported that SSNA1 filaments bind longitudinally along stabilized microtubules, induce microtubule branching and promote microtubule nucleation (Basnet et al. 2018). Growing microtubules undergo dynamic instability; a phenomenon whereby individual microtubules alternate between phases of growth and shrinkage via the transitions referred to as catastrophe and rescue (Mitchison and Kirschner 1984). Given its direct interaction with microtubules and localization to sites of dynamic microtubule growth in cells, SSNA1 is well-positioned to impact microtubule dynamics. Nonetheless, whether SSNA1 regulates dynamically-growing microtubules has not been investigated.

Microtubule regulation is not restricted to the dynamic microtubule ends. For example, microtubule-severing enzymes, motor proteins and mechanical forces induce microtubule lattice damage.The damaged microtubule lattice can be recognized and stabilized by MAPs (Schaedel et al. 2015, Aumeier et al. 2016, de Forges et al. 2016, Triclin et al. 2018, Vemu et al. 2018, Schaedel et al. 2019, Aher et al. 2020, Thery and Blanchoin 2020). SSNA1 has been identified as a binding partner of spastin, a microtubule-severing enzyme (Errico et al. 2004); and both SSNA1 and spastin localize at spindle poles and neuronal branch points (Yu et al. 2008, Goyal et al. 2014, Basnet et al. 2018). However, whether SSNA1 recognizes and stabilizes the damaged microtubule lattice is an open question.

In this study, we employed *in vitro* reconstitution techniques with purified protein components and TIRF microscopy to interrogate the roles of SSNA1 in regulating dynamic microtubule ends and lattices.

## RESULTS

### Human SSNA1 suppresses microtubule dynamicity

In order to investigate the effects of SSNA1 on dynamic microtubules, we purified human SSNA1 protein (Figure S1A-B) and employed an established TIRF-based microtubule dynamics *in vitro* reconstitution assay (Gell et al. 2010). Previous work implicated SSNA1 in spontaneous microtubule nucleation (Basnet et al. 2018). To build upon these observations, we assessed the ability of SSNA1 to promote templated microtubule nucleation from GMPCPP-stabilized microtubule seeds, thought to better reflect microtubule nucleation in cells (Wieczorek et al. 2015) (Figure 1A). We performed a titration of soluble tubulin in the presence and absence of 2.5 µM SSNA1 and found that SSNA1 promoted templated nucleation compared to microtubules grown with tubulin alone, thus supporting the role of SSNA1 in microtubule nucleation.

**Figure 1.**
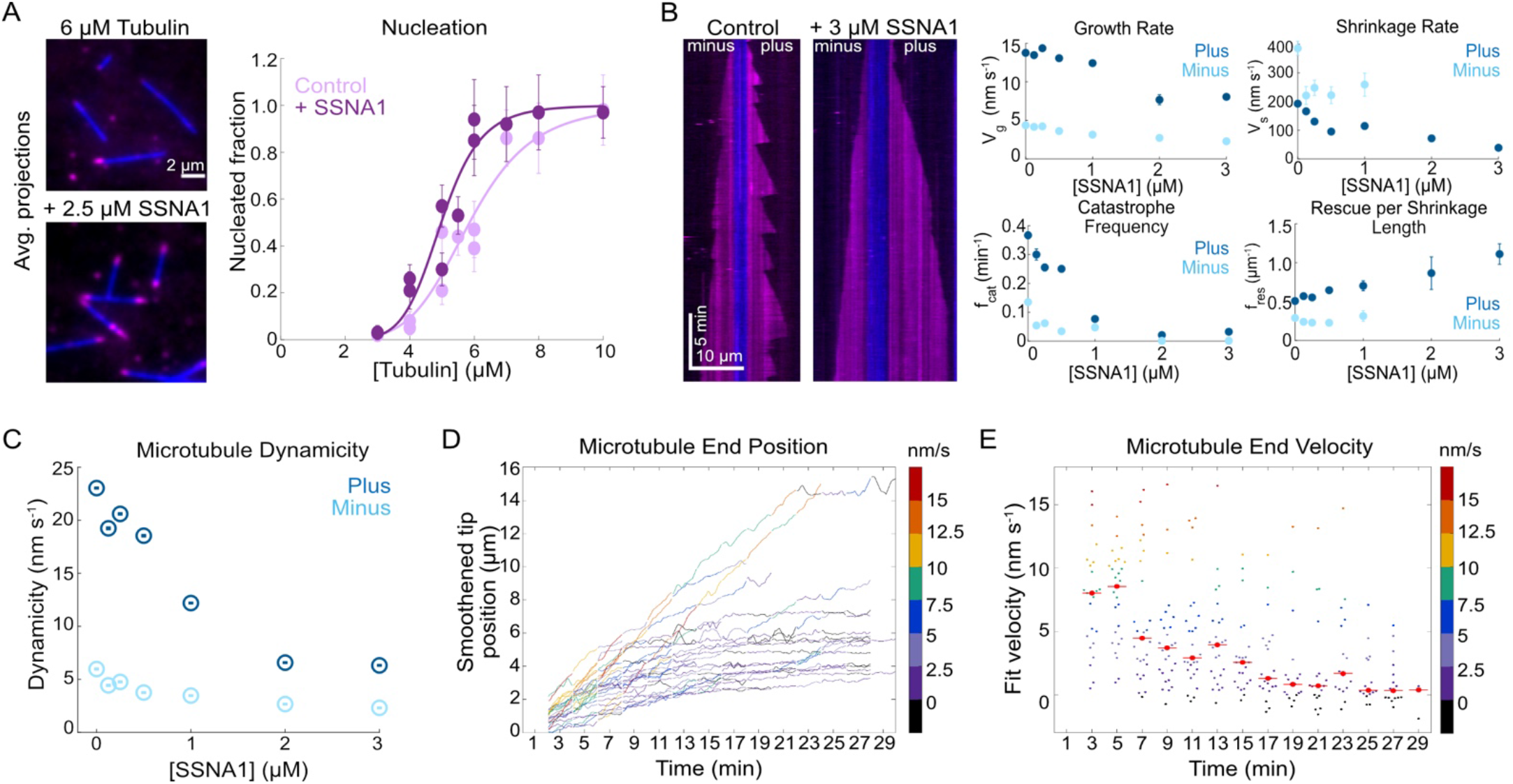
Human SSNA1 suppresses microtubule dynamicity and promotes microtubule nucleation. (A) Representative images of seeds (blue) and nucleated extensions (magenta) for the 6 µM tubulin condition with and without 2.5 µM 488-SSNA1 (left). Quantification of the fraction of seeds that nucleated in 15 minutes with tubulin alone and 2.5 µM 488-SSNA1 as a function of tubulin concentration (right). Data are individual experimental replicates ± SE from six experimental days (N = 30 - 68 microtubules for each concentration tested in the tubulin-alone control condition, N = 33 - 77 microtubules for each concentration tested in the SSNA1 condition). The data were fitted to a sigmoidal curve of the form *y*(*x*) = *x*s/(*C* + *x*s) (solid lines). For tubulin alone, C = 5.9 µM (95% CI: 5.5, 6.2) and s = 6.0 (95% CI: 3.4, 8.6). For the SSNA1 condition, C = 5.0 µM (95% CI: 4.7, 5.3) and s = 8.0 (95% CI: 4.3, 11.9). (B) Representative kymographs of microtubules grown with 9 µM Alexa-647 tubulin alone (control) and in the presence of 3 µM SSNA1; and quantification of the growth rate, catastrophe frequency, shrinkage rate and rescue per shrinkage length at the plus and minus ends of microtubules grown with 9 µM tubulin and concentrations of SSNA1 from 0 µM to 3 µM. Data are weighted means ± SE obtained from four independent experimental days (N = 43 - 485 growth events for each concentration tested at microtubule plus ends; N = 30 - 120 growth events for each concentration tested at microtubule minus ends). Note that no catastrophes were detected at the minus ends with SSNA1 concentrations greater than 1 µM, therefore, the shrinkage rate and rescue per shrinkage length could not be measured in those conditions. (C) Quantification of microtubule dynamicity as a function of the SSNA1 concentration. Dynamicity is calculated as total length of growth and shrinkage over the observation time. Data are weighted means ± SE. (D) Microtubule end positions as a function of time for microtubules grown with 9 µM tubulin and 3 µM SSNA1, color-coded with 2-minute segment velocity. Total of 27 microtubules were analyzed. (E) Microtubule end velocity in a 2-minute segment as a function of time calculated from the end positions in (D) and color-coded with velocity. For each bin, the median is shown as a bright red point with horizontal line.

Next, we performed SSNA1 titration experiments in which microtubules were grown with a range of SSNA1 concentrations, and quantified microtubule dynamics (Figure 1B, Videos 1-2). We found that SSNA1 suppressed the mean microtubule growth and shrinkage rates, suppressed catastrophe frequency and increased rescue frequency. The combined suppression of growth, shrinkage and catastrophe and promotion of rescue suggested that, overall, microtubules grown in the presence of SSNA1 were less dynamic. As such, we calculated microtubule dynamicity (defined as the total length of growth and shrinkage divided by the total time spent in growth and shrinkage) as a function of SSNA1 concentration. We found that microtubule dynamicity decreased with increasing SSNA1 concentrations at both plus and minus ends (Figure 1C).

Interestingly, we observed that the effects of SSNA1 on microtubule dynamics appeared to change over time. Notably, microtubule growth rate appeared to progressively slow down when microtubules were grown in the presence of SSNA1 (see kymograph in Figure 1B). To investigate this further, we tracked the ends of microtubules grown in the presence of 3 µM SSNA1 for up to 30 minutes (Figure 1D) and calculated the velocity of the microtubule end over time (Figure 1E). Quantification of the microtubule end velocity revealed a statistically-significant slow-down in the mean microtubule velocity (from 7.6 nm/s ± 0.9 nm/s at 3 min to 4.4 nm/s ± 0.8 nm/s at 9 min, p = 0.01, unpaired t-test); although, some microtubules did not slow down at all and others slowed down at later timepoints. Therefore, the microtubule growth rate progressively slows in the presence of SSNA1, but the extent and onset of the slowdown varies between individual microtubules.

### Progressive accumulation of SSNA1 on dynamic microtubules leads to a slowdown in microtubule growth and promotes microtubule end curvature

We hypothesized that the slow-down in microtubule growth rate over time in the presence of SSNA1 could be due to the progressive accumulation of SSNA1 on microtubules. To investigate SSNA1 localization on microtubules, we chemically labeled purified SSNA1 with Alexa-488 and Alexa-647 dyes. We first confirmed that the purified and labeled SSNA1 protein was able to self-assemble into filaments, as previously reported in the literature (Rodriguez-Rodriguez et al. 2011, Basnet et al. 2018) (Figure S1C). Next, we assessed the binding of labeled SSNA1 to GMPCPP-stabilized microtubules and found that SSNA1 progressively localized to the stabilized microtubules (Figure S2A). Quantification revealed a linear increase in SSNA1 fluorescence intensity on the microtubule lattice over time until approximately 5 minutes when the SSNA1 signal saturated. We also quantified the single-molecule dwell time of 488-SSNA1 on GMPCPP-stabilized microtubules and found that many SSNA1 molecules bound to the microtubule with durations in the seconds-range (Figure S2B). Thus, the slow build-up of SSNA1 on microtubules can be explained by the long single-molecule dwell times.

Next, we probed SSNA1 localization on dynamic microtubules and observed that SSNA1 intensity also increased over time on the dynamic microtubules (Figure 2A, Figure S3A, Video 3). Our quantitative analysis revealed that SSNA1 intensity measured at the microtubule end region inversely correlated with the microtubule end velocity (Figure 2B, Figure S3B-C). In addition, we noticed that the build-up of SSNA1 on dynamic microtubules promoted microtubule end curvature (Figure 2C). To investigate whether SSNA1 recognizes regions of increased microtubule curvature, we observed its localization on Taxol-stabilized microtubules, which display a broader range of curvatures (Figure S4). We found that SSNA1 does not specifically localize to curved microtubule regions, and therefore likely directly induces curvature of the dynamically growing microtubule ends.

**Figure 2.**
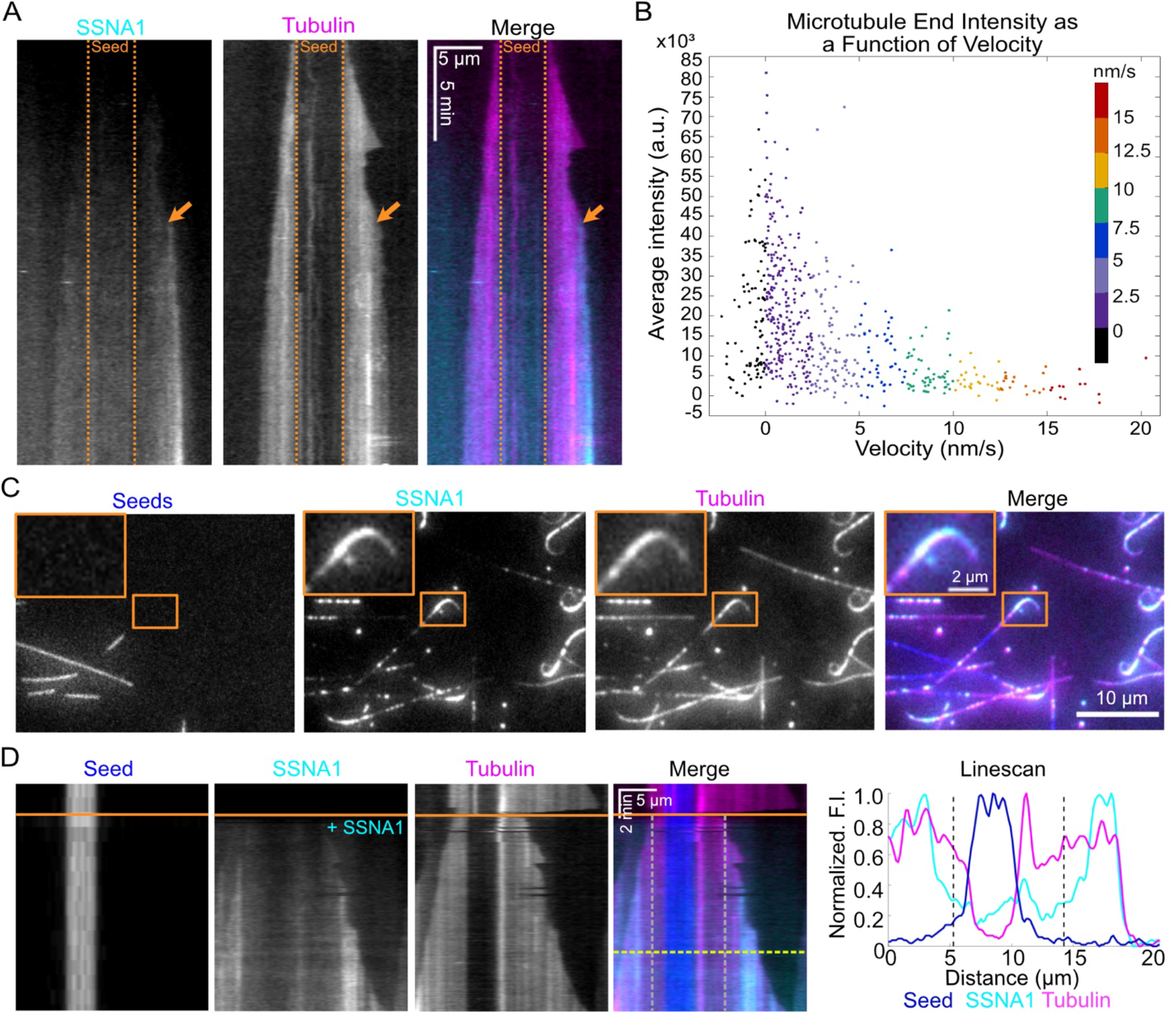
The progressive accumulation of SSNA1 on dynamic microtubules correlates with a slow-down in microtubule growth and promotes end curvature. (A) Kymograph of a microtubule grown with 9 µM Alexa-647-labeled tubulin and 5 µM Alexa-488-labeled SSNA1 (7% labeled) showing the progressive localization of 488-SSNA1 to a growing microtubule extension over time. The orange arrow indicates SSNA1 enrichment at the microtubule end. The dotted orange lines demarcate the position of the microtubule seed (not shown). (B) Plots of microtubule end velocities and corresponding SSNA1 fluorescence intensities at the microtubule end over time. Total of 26 microtubules were analyzed. (C) Representative images of microtubules grown with 9 µM Alexa-647-labeled tubulin and 5 µM Alexa-488-labeled SSNA1 (7% labeled) showing that SSNA1 promotes microtubule curvature. The orange box indicates the zoomed region shown in inset. (D) (Left) Dynamic microtubule extensions were pre-grown with 15 µM Alexa-647-tubulin, then 15 µM tubulin and 2.5 µM Alexa-488-SSNA1 (15% labeled) were introduced into the channel. The solid orange line indicates the time of SSNA1 introduction. The two dashed vertical lines mark the region of the microtubule grown without SSNA1. (Right) Linescan showing the normalized fluorescence intensities of the microtubule seed (blue), the dynamic microtubule extension (magenta) and SSNA1 (cyan); the dashed horizontal yellow line on the kymograph is the position of the linescan. The two dashed vertical lines mark the region of the microtubule grown without SSNA1, as denoted on the kymograph.

Our observation of SSNA1 localization revealed that SSNA1 does not bind uniformly along the entire microtubule lattice (Figure 2A, C). To investigate SSNA1’s binding preference to specific microtubule lattice regions, we first pre-grew dynamic microtubule extensions with 15 µM tubulin from GMPCPP-stabilized seeds in the absence of SSNA1 and then introduced 15 µM tubulin and 2.5 µM 488-SSNA1 into the reaction mix. This experimental set up allowed us to determine whether SSNA1 preferred to bind to GMPCPP-stabilized microtubule seeds; pre-existing, dynamically-grown microtubule lattices; or microtubule lattices that were grown in the presence of SSNA1. Interestingly, we found that SSNA1 preferentially bound to the newly-growing extensions, as opposed to the pre-existing parts of the microtubule lattice (Figure 2D, Video 4). Taken together, our results show that SSNA1 initially loads onto the dynamic end, and subsequently persists along the newly-polymerized microtubule lattice, forming stretches of lattice accumulation.

### SSNA1 protects dynamic microtubules against induced catastrophe

Thus far, our experiments demonstrated that SSNA1 accumulates in stretches at microtubule ends, where it suppresses microtubule catastrophe. In cells, microtubules are frequently subject to specific destabilizing perturbations; we therefore decided to further investigate the ability of SSNA1 to protect microtubules against induced catastrophe. First, we tested whether SSNA1 could protect microtubules from catastrophe induced by dilution of soluble tubulin. Specifically, microtubules were grown with 9 µM tubulin in the presence or absence of 5 µM SSNA1 for 30 minutes and then the reaction mixture was exchanged for assay buffer, effectively removing all soluble tubulin and SSNA1 protein from solution (Figure 3A, Figure S5A, Videos 5 & 6). We quantified the rate of polymer loss after tubulin dilution and found that control microtubules lost polymer at a rate of 24 nm s^-1^ ± 2 nm s^-1^ (N = 141, 3 experimental repeats) while microtubules that were grown with SSNA1 lost polymer at a rate of 0.8 nm s^-1^ ± 0.4 nm s^-1^ (N = 181, 3 experimental repeats) (Figure 3A). Dilution experiments using fluorescent SSNA1 revealed that SSNA1 stayed bound to the microtubule after dilution, protecting the microtubule end from catastrophe (Figure S5B).

**Figure 3.**
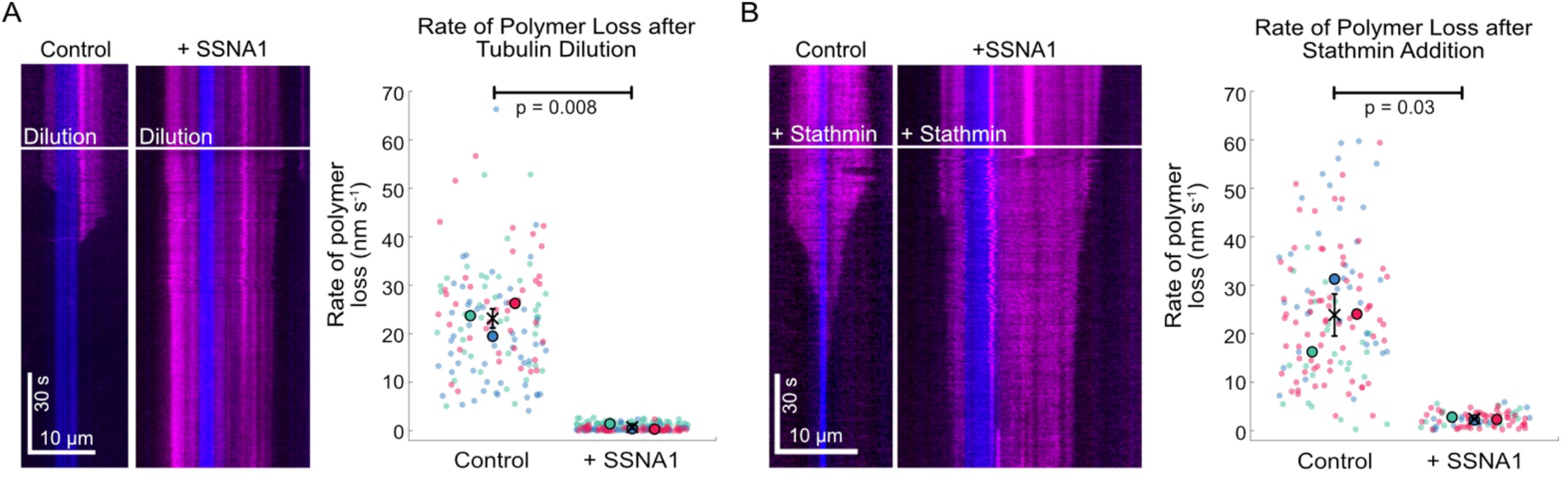
SSNA1 protects microtubule against catastrophe induced by either tubulin dilution or stathmin-mediated tubulin sequestration. (A) Microtubules were grown with 9 µM tubulin ± 5 µM SSNA1 for 30 min. After 30 minutes, the buffer was exchanged for pre-warmed assay buffer containing no soluble tubulin. The white lines on the kymographs indicate the time of tubulin dilution. Quantification of the rate of polymer loss after dilution is shown in the graph on the right (N = 141 for control, N = 181 for SSNA1, both over 3 experimental repeats). (B) Experiments were performed as in (A) except the reaction mixture was exchanged for 9 µM tubulin ± 6 µM stathmin. The white lines on the kymographs indicate the time of reaction exchange. Quantification of the rate of polymer loss after buffer exchange is shown in the graph on the right (N = 140 for control, N = 85 for SSNA1, both over 3 experimental repeats). Data in (A) and (B) are plotted in “SuperPlots” (see also methods) and statistical significance was determined by Welch’s t-test on the means of each experimental replicate. The colors of the individual data points represent experimental days, and the means are plotted as larger points in the same color. The cross indicates the average of the experiment means and the vertical bar is standard error of the mean.

One way in which the soluble tubulin concentration is regulated in cells is through the sequestration of tubulin dimers by members of the Op18/stathmin family proteins (Belmont and Mitchison 1996, Cassimeris 2002, Steinmetz 2007, Gupta et al. 2013). Therefore, we next tested whether SSNA1 could also protect microtubules against stathmin activity (Figure 3B). As before, microtubules were grown with 9 µM tubulin in the presence or absence of 5 µM SSNA1 for 30 minutes, but the reaction mixture was exchanged for 9 µM tubulin and 6 µM stathmin. We found that control microtubules lost polymer at a rate of 25 nm s^-1^ ± 4 nm s^-1^ (N = 140, 3 experimental repeats) while microtubules that were grown with SSNA1 lost polymer at a rate of 2.7 nm s^-1^ ± 0.2 nm s^-1^ (N = 85, 3 experimental repeats) in the presence of stathmin. Exchanging the reaction mixture for 9 µM tubulin alone did not cause catastrophe (Figure S5C). Taken together, these experiments reveal that SSNA1 protects microtubules from catastrophe even in the absence of available soluble tubulin.

### SSNA1 inhibits spastin-mediated microtubule severing and recognizes sites of lattice damage

In addition to the factors that induce microtubule catastrophe, microtubules in cells are also destabilized by the action of microtubule-severing enzymes, which play important roles in regulating the dynamics and organization of microtubule networks (McNally and Roll-Mecak 2018, Kuo and Howard 2021). We therefore asked whether SSNA1 could protect microtubules from the action of spastin, a microtubule-severing enzyme. To test whether SSNA1 protects microtubules against severing, we incubated GMPCPP-stabilized microtubules with or without 1 µM 488-SSNA1 for 10 minutes and then introduced 200 nM spastin (Figure 4A, Video 7). We found that, while spastin was able to efficiently sever control microtubules, no severing was observed for the SSNA1-coated microtubules over 30 minutes. We next tested whether SSNA1 recognizes sites of microtubule lattice damage by incubating microtubules with 100 nM spastin for 5 minutes to induce damage and then exchanging the solution to 5 µM 647-SSNA1 alone (Figure 4B). We observed that SSNA1 preferentially bound to regions of the microtubule lattice with lower fluorescence intensity, characteristic of sites of lattice damage. Linescan analysis of the SSNA1 and microtubule fluorescence intensities revealed that SSNA1 progressively built up at the sites of microtubule damage over the course of the experiment (Figure 4B, Figure S6A, Video 8). In addition, we found that SSNA1 recognized sites of lower tubulin intensity (i.e., lattice defect sites) on Taxol-stabilized microtubules (Figure S6B). Strikingly, we were also able to detect SSNA1 enrichment on dynamically growing microtubule lattices in regions of lower tubulin intensity (Figure 4C). Therefore, SSNA1 is a specific sensor of naturally-occurring and spastin-induced microtubule lattice damage.

**Figure 4.**
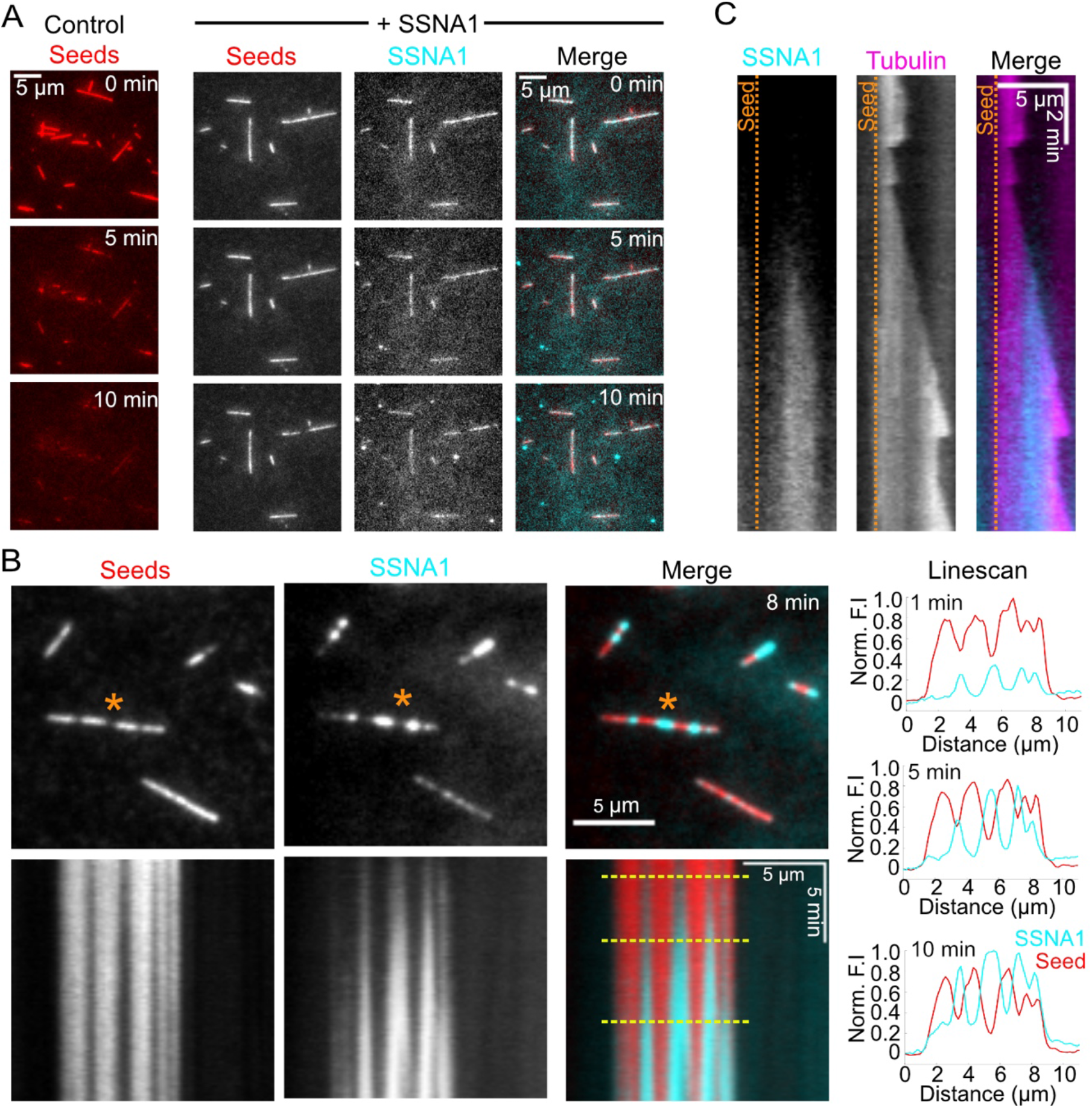
SSNA1 inhibits microtubule severing by spastin and recognizes sites of microtubule lattice damage. (A) Representative images of control (left) and SSNA1-coated (right) microtubules incubated with 200 nM human spastin. Microtubules were pre-incubated with either SSNA1 storage buffer (control) or 1 µM 488-SSNA1 for 10 minutes prior to the addition of spastin. Spastin was introduced at t = 0 min. (B) Representative images of GMPCPP-stabilized microtubules that were pre-incubated for 5 minutes with 100 nM spastin and then subsequently incubated with 5 µM 647-SSNA1. Images are from 8 minutes after SSNA1 addition. Below are kymographs showing SSNA1 localization to sites of spastin-induced microtubule damage. The orange asterisks indicate the microtubule used for kymograph and linescan analysis. The dashed yellow lines indicate the position of the linescans (right) of the seed and SSNA1 fluorescence intensities at 1 min, 5 min and 10 min after the introduction of SSNA1. (C) Kymograph showing 488-SSNA1 enrichment at a site of microtubule lattice damage on a dynamically growing microtubule extension. Microtubules were grown with 9 µM Alexa-647-labeled tubulin and 5 µM Alexa-488-labeled SSNA1 (7% labeled). The dotted orange line demarcates the position of the microtubule seed (not shown).

## DISCUSSION

SSNA1 plays important roles in several fundamental, microtubule-based cellular processes including cilia formation, cell division and axonal branching (Pfannenschmid et al. 2003, Schoppmeier et al. 2005, Lai et al. 2011, Goyal et al. 2014, Basnet et al. 2018). Despite an appreciation of the biological functions of SSNA1, an understanding of the direct effects on dynamic microtubules remained lacking. In this study, using *in vitro* reconstitution approaches, we explored the effects of SSNA1 on microtubules and found that SSNA1 robustly stabilizes dynamic microtubules and detects sites of lattice damage, occurring both naturally and induced by spastin.

Microtubule-stabilizing proteins have critical functions in regulating microtubules in cells. One of the best studied classical stabilizing MAPs is tau, known for its roles in neurons and involvement in neurodegenerative diseases (Morris et al. 2011, Iqbal et al. 2016, Gao et al. 2018, Barbier et al. 2019). Studies in cells and *in vitro* report that tau forms oligomers on the outer microtubule surface (Al-Bassam et al. 2002), and stabilizes microtubules against depolymerization (Drechsel et al. 1992, Ramirez-Rios et al. 2016, Prezel et al. 2018). Similarly, SSNA1 forms filaments that bind longitudinally on microtubules (Rodriguez-Rodriguez et al. 2011, Basnet et al. 2018). Our work demonstrates that SSNA1 simultaneously modulates all four parameters of microtubule dynamic instability – slowing down the rates of growth, shrinkage and catastrophe, and promoting rescue. Remarkably, we find that SSNA1 prevents catastrophe even when all tubulin is removed from solution or sequestered by a microtubule destabilizer Op18/Stathmin. Thus, SSNA1 is a potent microtubule stabilizing protein and this activity likely underlies its cellular function.

In contrast to tau, which has been reported to promote microtubule growth (Drechsel et al. 1992), the growth rate slows down in the presence of SSNA1. One way to slow down microtubule growth is through sequestration of soluble tubulin – this is the mechanism employed by Op18/Stathmin (Belmont and Mitchison 1996, Arnal et al. 2000, Cassimeris 2002, Steinmetz 2007, Gupta et al. 2013). However, tubulin sequestration would affect the growth rate of all microtubules simultaneously; this is not the case with SSNA1, where the onset of slow-down occurs at different times for individual microtubules. Instead, we find that suppression of microtubule growth rate correlates with the progressive SSNA1 accumulation on microtubule ends over time. A slow-down in growth is typically observed just prior to microtubule catastrophe. Microtubule growth slow-down may be a consequence of perturbations in the dynamic end structure, including incomplete tubules, exposed protofilaments or ragged microtubule ends. Notably, previous data from cryo-ET experiments indicate that SSNA1 can bind to partial tubule structures, as it supports the growth of protofilaments away from the mother microtubule (Basnet et al. 2018). It is thus possible that SSNA1 recognizes specific dynamic end structures that are incompatible with continued unperturbed growth.

It has recently been proposed that taxanes, microtubule-stabilizing compounds widely used in cancer therapy, recognize microtubule ends that are in a pre-catastrophe state (Rai et al. 2020). Significantly, the behavior of SSNA1 on dynamic microtubules is highly reminiscent of taxanes: at sub-saturating concentrations, taxanes accumulate at growing microtubule ends following growth perturbations, and form persistent patches that stabilize the microtubule lattice (Rai et al. 2020). While taxanes show a preference for GMPCPP-grown parts of the microtubule lattice, which mimic the extended GTP-tubulin conformation, we do not see enhanced localization of SSNA1 on the GMPCPP-lattice regions. Therefore, we do not think that nucleotide state recognition is the primary mechanism for SSNA1 localization. Electron microscopy data demonstrate that taxanes accumulate on incomplete microtubule lattice structures (Rai et al. 2020). Similarly, we find that patches of SSNA1 accumulate on dimmer regions of dynamic microtubule lattices (i.e., incomplete tubules) and growing microtubule ends. Thus, our data support a mechanism in which SSNA1 binding is determined by the structure of the microtubule lattice. Furthermore, we demonstrate that SSNA1 detects sites of spastin-mediated lattice damage. Hence, like taxanes, SSNA1 senses microtubule lattice damage.

Several lines of evidence support an important functional interplay between SSNA1 and spastin: SSNA1 is a binding partner of spastin (Errico et al. 2004); SSNA1 and spastin colocalize in dividing cells (Goyal et al. 2014); SSNA1 and spastin promote axonal branching in developing neurons (Yu et al. 2008, Goyal et al. 2014, Basnet et al. 2018); and SSNA1’s putative spastin-binding domain is required to promote axonal branching (Goyal et al. 2014). Our data demonstrate that SSNA1 can both inhibit spastin activity and detect spastin-induced damage, further highlighting the interplay between the two proteins. Similarly, Tau condensates on microtubule lattices were recently found to protect against severing by both spastin and katanin (Siahaan et al. 2019, Tan et al. 2019). SSNA1’s microtubule stabilizing activity is similar to that of CAMSAP2 and CAMSAP3: minus-end stabilizing proteins that associate with new minus ends and form stretches of stabilized lattice (Jiang et al., 2014). Interestingly, katanin interacts with CAMSAP2 and CAMSAP3 and limits the length of CAMSAP2-stretches on the microtubule minus ends. Thus, balancing the activity of microtubule severing and stabilizing proteins may represent a general mechanism for regulating microtubule number and mass in cells. Perhaps counterintuitively, spastin is implicated in microtubule network amplification by generating new microtubules fragments after severing (Vemu et al. 2018, Kuo et al. 2019, Kuo et al. 2019, Kuo and Howard 2021). Whether synergy between SSNA1’s stabilizing activity and spastin’s severing activity is critical for microtubule network amplification in cells presents an important area for future work.

Microtubule dynamics and network organization are influenced by the properties of both the microtubule end and lattice. Herein, we have identified SSNA1 as both a microtubule stabilizing protein and sensor of microtubule damage. Our work provides mechanistic insight into an important but understudied microtubule regulatory protein with essential functions in cilia, cell division and developing neurons.

## MATERIALS AND METHODS

### DNA constructs

The cDNA encoding human SSNA1 (NM_003731.2) with an N-terminal 6xHis-tag in a pReceiver-B01 vector was purchased from GeneCopoeia, Rockville, MD, USA (product ID: Q0661). The cDNA encoding human stathmin/Op18 with an N-terminal 6xHis-tag in a pET15b vector was a kind gift from the Goodson lab (University of Notre Dame, USA) (Gupta et al., 2013). The cDNA encoding human spastin (NM_014946.3) was purchased from GeneCopoeia (product ID: U1177) and was subcloned into a modified pET vector containing N-terminal 6xHis and MBP tags. The pET MBP His6 LIC cloning vector (2Cc-T) was a gift from Scott Gradia (Addgene plasmid # 37237; http://n2t.net/addgene:37237; RRID: Addgene_37237).

### Protein preparation

Bovine brain tubulin was purified using cycles of polymerization and depolymerization using the high-molarity PIPES method (Castoldi and Popov 2003). Tubulin was labeled with tetramethylrhodamine (TAMRA), Alexa Fluor 488 and Alexa Fluor 647 dyes (ThermoFisher Scientific, Waltham, MA, USA) according to the standard protocols and as previously described (Hyman et al. 1991, Gell et al. 2010). Fluorescently-labeled tubulin was used at a ratio of between 5% and 10% of the total tubulin.

Human 6His-SSNA1 was expressed in BL21 DE3 Gold cells in Studier autoinduction media (Teknova, Hollister, CA, SUA; cat. #3S2000) for 96H. Expression cell pellets were lysed for 1 hr at 4°C in 50 mM HEPES (pH 7.5), 150 mM NaCl, 10% (v/v) glycerol, 10 mM imidazole and 1 mM DTT and supplemented with 1 mg/ml lysozyme, 10 mg/ml PMSF, EDTA-free protease inhibitors (Roche, Basel, Switzerland), and 25 U/ml Pierce universal nuclease (Invitrogen). The crude lysate was sonicated on ice and clarified by centrifugation for 30 min at 4°C and 35,000 rpm in a Beckman L90K Optima and 50.2 Ti rotor (Beckman, Brea, CA, USA). The clarified lysate was applied to a HisTrapHP column (Cytiva, Marlborough, MA) according to the manufacturer’s protocol and eluted with 50 mM HEPES (pH 7.5), 150 mM NaCl, 10% (v/v) glycerol, 1 mM DTT and a linear gradient of 50 - 500 mM imidazole. The eluted protein was buffer exchanged using a PD-10 desalting column (Cytiva) into 20 mM HEPES (pH 7.5), 150 mM NaCl, 10% (v/v) glycerol and 1 mM DTT. Purified SSNA1 was labeled using Alexa Fluor 488 and Alexa Fluor 647 Microscale Protein Labeling Kits (ThermoFisher Scientific, cat. #A30006 and #A30009) according to the manufacturer’s instructions. Protein purity was assessed by SDS-PAGE and mass spectrometry analysis.

The spastin purification protocol was adapted from (Kuo et al. 2019). Human 6His-MBP-spastin was expressed in Rosetta(DE3) cells. Expression was induced with 0.5 mM IPTG and expressed overnight at 16°C. Cells were lysed for 1H at 4°C in 30 mM HEPES (pH 7.4), 300 mM NaCl, 10 mM imidazole, 5% glycerol, 2 mM DTT, 10 µM ATP and 2 mM DTT and supplemented with 1 mg/ml lysozyme, 10 mg/ml PMSF and EDTA-free protease inhibitors. The crude lysate was sonicated on ice and then clarified by centrifugation for 30 min at 4°C and 35,000 rpm in a Beckman L90K Optima and 50.2 Ti rotor. Clarified lysates were applied to a HisTrapHP column (Cytiva) according to the manufacturer’s protocol. His-tagged protein was eluted with 30 mM HEPES (pH 7.4), 300 mM NaCl, 10 µM ATP, 5% (v/v) glycerol and 2 mM DTT and linear gradient of 50 mM - 500 mM imidazole. For storage, the buffer was exchanged to 20mM HEPES (pH 7.4), 0.15 M NaCl, 5% glycerol, 0.5 mM DTT, 10 µM ATP and PMSF using PD-10 desalting columns (Cytiva).

Human 6His-stathmin/Op18 protein was expressed in E. *coli* and purified as previously described (Arnal et al. 2000). All proteins were snap frozen in liquid nitrogen and stored at -80°C.

### Imaging assay

All imaging was performed using a Nikon Eclipse Ti microscope with a 100×/1.49 n.a. TIRF objective (Nikon, Tokyo, Japan), Andor iXon Ultra EM-CCD (electron multiplying charge-coupled device) camera (Andor, Belfast, UK); 488-, 561-, and 640-nm solid-state lasers (Nikon Lu-NA); HS-625 high speed emission filter wheel (Finger Lakes Instrumentation, Lima, NY USA); and standard filter sets. An objective heater was used to maintain the sample at 35°C. Microscope chambers were constructed as previously described (Gell et al. 2010). Briefly, 22 × 22 mm and 18 × 18 mm silanized coverslips were separated by strips of Parafilm to create narrow channels for the exchange of solution. Images were acquired using NIS-Elements (Nikon) with exposure times of 50 ms – 100 ms and at the frame rates specified in the methods.

### Microtubule dynamics

For microtubule dynamics experiments, GMPCPP-stabilized microtubules seeds were prepared according to standard protocols (Gell et al. 2010, Chen and Doxsey 2012). Dynamic microtubule extensions were polymerized from surface-immobilized GMPCPP-stabilized templates as described previously (Gell et al. 2010). Imaging buffer containing soluble tubulin ranging from 9 µM tubulin, 1 mM GTP, and proteins at the concentrations indicated in the text were introduced into the imaging chamber. The imaging buffer consisted of BRB80 supplemented with 40 mM glucose, 40 µg/ml glucose oxidase, 16 µg/ml catalase, 0.5 mg/ml casein, 50 mM KCl, 10 mM DTT and 0.1% methylcellulose. Dynamic microtubules were grown with or without unlabeled SSNA1 and imaged for 30 minutes (0.2 fps).

Quantification of microtubule dynamics parameters was performed using kymographs generated in Fiji (Schindelin et al. 2012) as described previously (Zanic 2016). Catastrophe frequency was calculated by dividing the number of catastrophes by the total time spent in the growth phase. Rescue was calculated by dividing the number of rescues observed by the total shrinkage length. The error for catastrophe frequency and rescue per shrinkage length are counting errors. Microtubule dynamicity was calculated by summing the total length of growth and shrinkage and dividing by the total observation time (Toso et al. 1993). The error for dynamicity was calculated as pixel size (160 nm) multiplied by √(N)/T where N is the number of points marked on the kymographs.

For the analysis of microtubule growth rate over time, microtubule end positions were determined using KymographClear and KymographDirect using tubulin channel (Mangeol et al. 2016). A custom MATLAB (The MathWorks, Natick, MA, USA) code was used to determine growth rate as a function of time. Briefly, a linear function was fit to position and time data points within a 2-minute window to determine the mean velocity. The window was then shifted to the next 2-minute interval and fitting procedure was repeated until the end of the trajectory. To focus on the characterization of microtubule growth over time, the segments with negative velocity at the beginning of a trajectory were eliminated (i.e., segment contains shrinkage phase), such that each trajectory starts with a positive velocity segment, while subsequent segments with negative velocity were kept. For each 2-minute segment, position data were color-coded based on the determined velocity and plotted as a function of time. For further analysis of growth segments, segments determined to have velocities smaller than -2.5nm/s were classified as shrinking segments, and were not included. Segments from multiple microtubule trajectories for a given time window were grouped and the median for each time window was calculated. Similarly, the velocity-versus-time data points were color-coded based on the segment velocity.

### Microtubule nucleation

The templated microtubule nucleation experiments were based on (Wieczorek et al. 2015): GMPCPP-stabilized seeds were incubated with tubulin concentrations ranging from 0 – 10 µM in the presence and absence of 2.5 µM SSNA1 and imaged every 15 seconds for 15 minutes. Nucleation was measured as the fraction of individual GMPCPP seeds that were observed to grow at least one microtubule extension of >3 pixels in length (480 nm) within 15 minutes. Analysis was performed on maximum projection images from the 15-minute time-lapse Videos. Data across the range of the tubulin concentrations were fitted to the sigmoidal equation *y*(*x*) = *x*s/(*C* + *x*s) in MATLAB, as previously published (Wieczorek et al. 2015), where C is the half maximal concentration at which nucleation occurs and s is the steepness of the curve. The errors were calculated as √N/total number seeds, where N is the number of nucleated seeds, except when 0 seeds nucleated, then the error was estimated as 1/total number of seeds.

### SSNA1 localization on Taxol-stabilized microtubules

Taxol-stabilized microtubules were prepared as follows: microtubules were grown with 32 µM tetra-rhodamine-labeled tubulin (25% labeled) for 30 minutes at 35°C and stabilized with 10 µM Taxol (Tocris, Minneapolis, MN, USA; Cat. # 1097) in BBR80 buffer. The taxol-stabilized microtubules were then spun in an airfuge for 5 minutes at 20 psi and sheared with an 18-gauge needle. Taxol microtubules were bound to the coverslip surface of a flow cell using anti-Rhodamine antibody, then 5µM 488-SSNA1 was introduced and the microtubules were imaged every 30 seconds for 60 minutes.

### SSNA1 localization on GMPCPP-stabilized microtubules

GMPCPP-stabilized microtubules adhered to coverslips were incubated with 2 µM Alexa-488-SSNA1 and imaged for 10 minutes (0.2 fps). The total SSNA1 fluorescence intensity along the total length of microtubule lattice in the microscope field of view excluding the background was measured in every frame (5 s interval) of the 10-minute movie using Fiji. The background from the SSNA1 channel was excluded by creating a mask around the microtubules, applying the mask to the SSNA1 channel, and including only the areas occupied by microtubules in the SSNA1 intensity measurements.

### SSNA1 localization on dynamic microtubules

For SSNA1 localization experiments on dynamic microtubules, microtubules were grown from GMPCPP-microtubule seeds with 9 µM tubulin and 5 µM 488-SSNA1 and imaged for 60 minutes (0.2 fps).

For the mixed-lattice wash-in experiments (Figure 2D), dynamic microtubules were pre-grown from GMPCPP-microtubule seeds with 15 µM tubulin for 15 minutes and then the reaction was exchanged for 15 µM tubulin and 2.5 µM Alexa-488-SSNA1 (15% labeled). The microtubules were imaged for 60 minutes in total (2 minutes prior to exchange and 58 minutes after exchange) at 0.2 fps.

To quantify SSNA1 localization on growing microtubules over time, the microtubule end velocity was determined for 2-minute segments, and the mean SSNA1 intensity at the microtubule end was determined using a custom MATLAB code. For each time frame (i.e., horizontal line on a kymograph) within 2-minute segment: i) mean solution background intensity was calculated within a 5-pixel-long region located 3 pixels away from the tracked end position, ii) mean lattice intensity was calculated within a 5-pixel-long region on the microtubule lattice ending at the tracked end position, iii) the mean solution background intensity was subtracted from mean lattice intensity to obtain mean SSNA1 intensity. Time frames which had <5 pixels available for background intensity calculation were eliminated. Next, an average SSNA1 intensity within 2-minute segment was calculated using single-time-frame (frame interval = 5 s) mean SSNA1 intensities. The average SSNA1 intensity data points were color-coded with 2-minute segment velocity and plotted as a function of time and velocity.

### Tubulin dilution/sequestration experiments

For the tubulin dilution and stathmin-mediated tubulin-sequestration experiments, microtubules were first grown with 9 µM 488-tubulin for 30 minutes in the absence and presence of 5 µM SSNA1 and then the reaction solution was manually exchanged using filter paper. For the dilution experiments, the reaction solution was exchanged to imaging buffer. For the stathmin experiments, the reaction solution was exchanged to 9 µM tubulin and 6 µM stathmin. Microtubules were imaged at 2 fps for 30 seconds prior to the exchange and for 10 minutes following the exchange. The mean rate of polymer loss was calculated from the microtubule end position at the time of solution exchange until the time the end depolymerized completely to the microtubule seed (for control microtubules) or over 5 minutes (for the SSNA1-grown microtubules that did not undergo catastrophe within 5 minutes). No distinction was made between microtubule plus and minus ends. Outliers were identified as the top and bottom 2 percentiles and discarded. Data were plotted as “SuperPlots” in which individual data points are color-coded by experiment (Lord et al. 2020) and statistical significance testing was performed on the means of the experimental repeats using Welch’s t-test.

### Single molecule dwell time analysis

GMPCPP-seeds were incubated with 5 nM 488-SSNA1 (40% labeled) and imaged for 10 minutes at 1 fps using maximum laser power and 100 ms exposure. The durations of the binding events were measured from kymographs and plotted as a histogram in MATLAB. The mean dwell time was determined by the decay constant of an exponential curve fitted to the data. The association rate was calculated as the number of binding events per second per nM per total length of the microtubule seeds in µm.

### Microtubule severing and damage assays

For all experiments using spastin, the imaging buffer was supplement with 1 mM ATP and 1 mM MgCl2 was included whenever spastin was used. For the microtubule severing assay, GMPCPP-stabilized microtubules were incubated with 1 µM 488-SSNA1 or SSNA1 storage buffer (control) for 10 minutes, then 200 nM spastin and 1 µM 488-SSNA1 was introduced into the flow cell and the microtubules were imaged for 30 minutes at 0.5 fps.

For the damage recognition assay, GMPCPP-stabilized microtubules were first incubated for 5 minutes with 100 nM spastin to generate microtubule lattice damage. The reaction was washed out with BRB80 and imaging buffer and then the damaged microtubules were incubated with 5 µM 647-SSNA1 and imaged every 10 seconds for 30 minutes.

## ACKNOWLEDGEMENTS

We thank S. Hall and J. Hayes for help with protein purification, C. Strothman for the MATLAB dynamics analysis script, and H. Goodson (University of Notre Dame). We thank H. McDonald and the Vanderbilt Mass Spectrometry Research Center (MSRC) Cores for the mass spectrometry analysis, which was supported in part by Vanderbilt Ingram Cancer Center Resource Share Scholarship 2020-2909607. We thank members of the Zanic lab and the Vanderbilt Microtubules and Motors Club for discussions and feedback. E.J.L. acknowledges the support of the National Institutes of Health IBSTO training grant T32CA119925. M.Z. acknowledges support from the National Institutes of Health grant R35GM119552, and the National Science Foundation grant MCB2018661.

The authors declare no competing interest.

## SUPPLEMENTAL FIGURES

**Figure S1.**
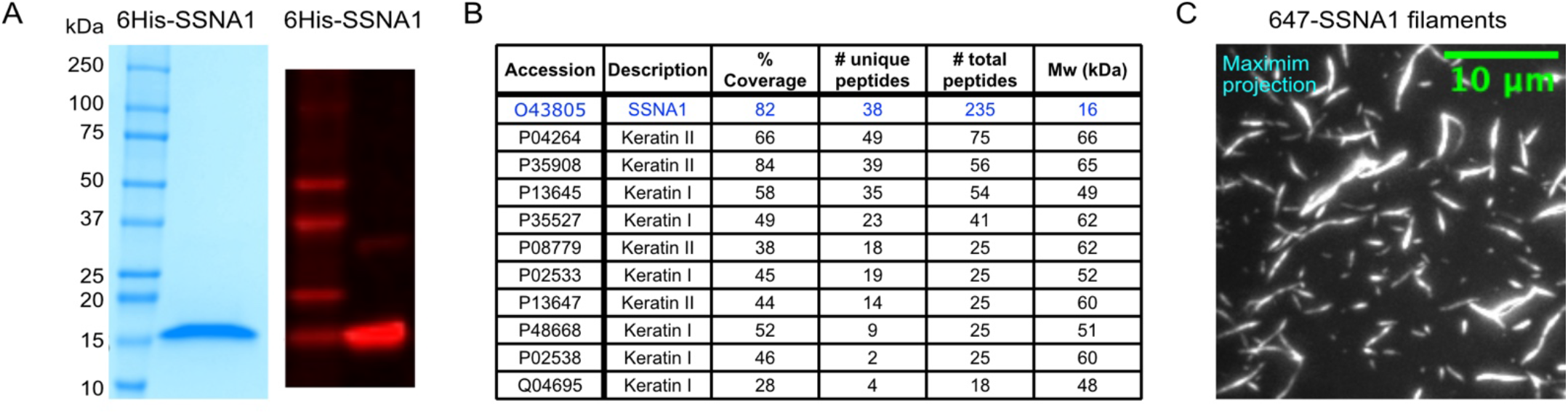
Purified human SSNA1 forms filaments. (A) SDS-page gel showing purified His-SSNA1 protein (left) and western blot using SSNA1 Rabbit Polyclonal Antibody (Proteintech; cat# 11797-1-AP). (B) Mass spectrometry analysis of His-SSNA1 purified from E. *coli* cells. The hits with more than 10 total peptides are listed. (C) 647-SSNA1 filaments were grown by incubating 58 µM 647-His-SSNA1 at 35°C for 4 hours. The reaction was then flowed onto coverslips coated with anti-His-antibody and imaged by TIRF microscopy. Image is a maximum projection image of multiple fields of view.

**Figure S2.**
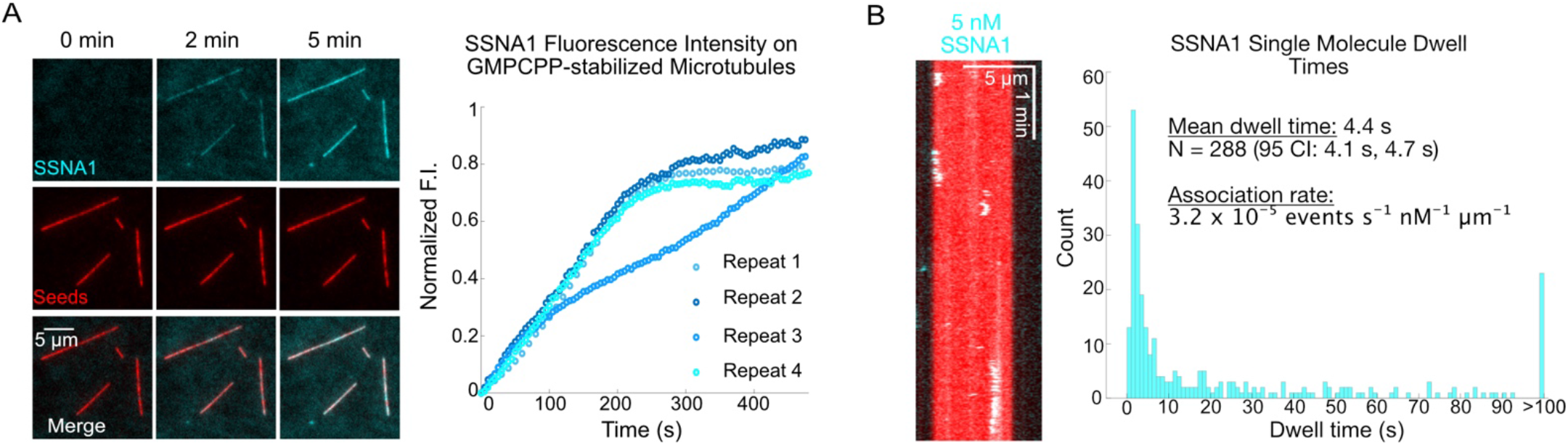
SSNA1 binds to GMPCPP-stabilized microtubules with a long single molecule dwell time. (A) Representative images showing GMPCPP-stabilized microtubules incubated with 2 µM Alexa-488-SSNA1. The total SSNA1 fluorescence intensity along the total length of microtubule lattice in the microscope field of view excluding the background was measured over time for 4 independent experimental repeats. (B) Quantification of 488-SSNA1 single-molecule dwell time on GMPCPP-stabilized microtubules that were incubated with 5 nM 488-SSNA1.

**Figure S3.**
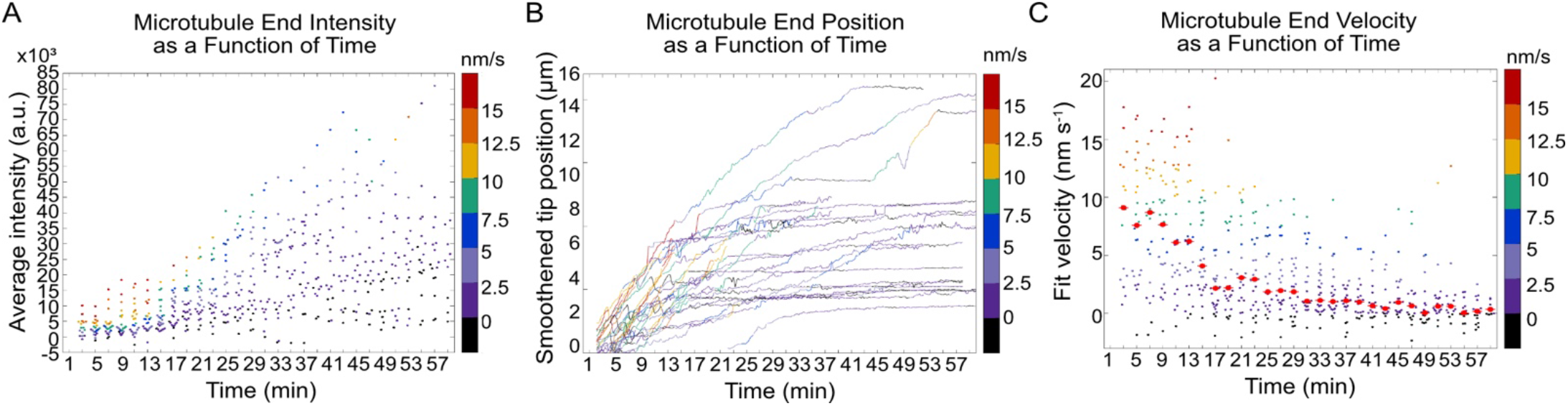
SSNA1 intensity on the microtubule end increases over time and end velocity decreases. (A) Average SSNA1 intensity at the microtubule end as a function of time, color-coded with 2-minute velocity. (B) Microtubule end positions as a function of time, color-coded with 2-minute segment velocity. (C) Microtubule end velocity in a 2-minute segment as a function of time, color-coded with velocity. For each bin, the median is shown as a bright red point with horizontal line.

**Figure S4.**
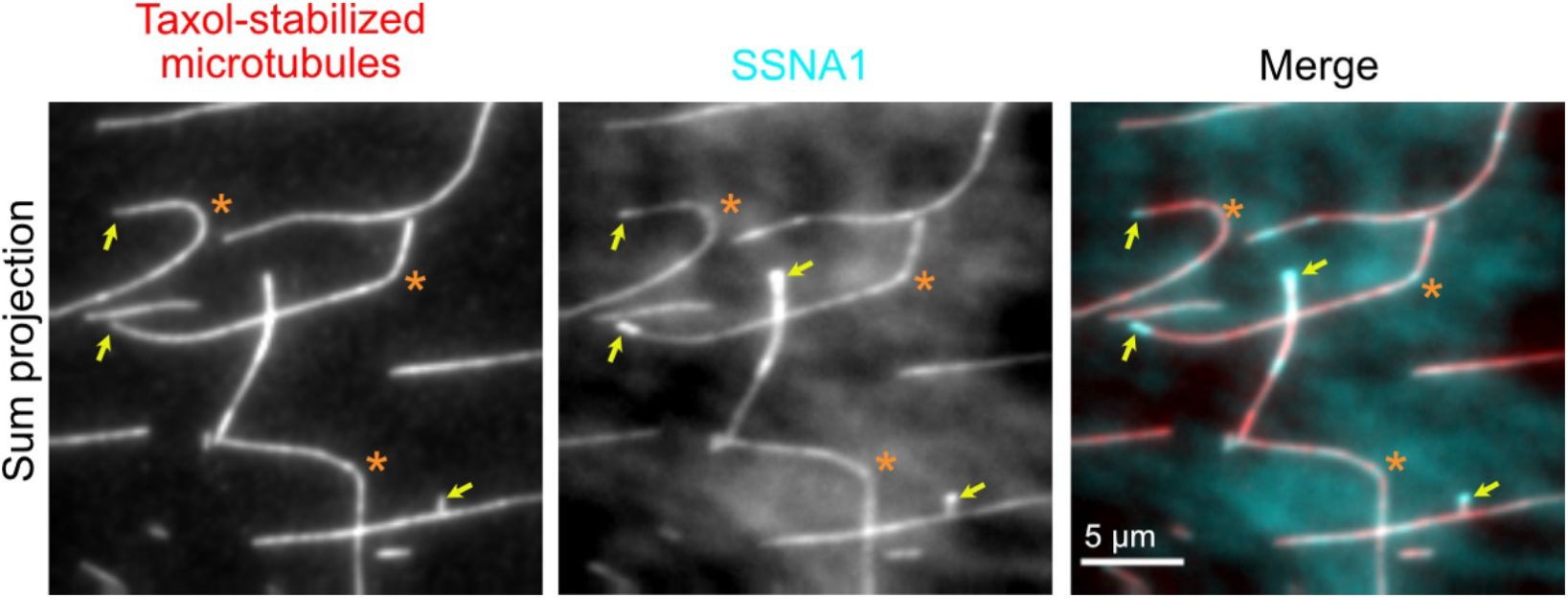
SSNA1 does not recognize microtubule curvature. Taxol-stabilized microtubules were incubated with 5µM 488-SSNA1 and imaged for 60 minutes. Sum projection images of a representative field of view are shown. Orange asterisks indicate curved microtubule regions. Yellow arrows indicate enhanced binding of SSNA1 to some microtubule ends.

**Figure S5.**
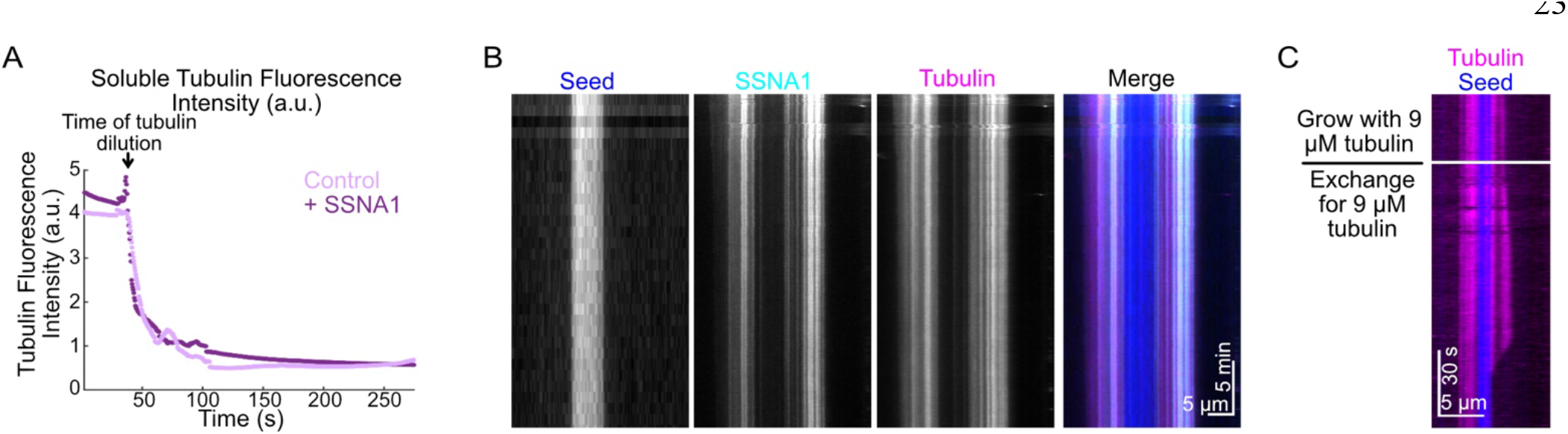
SSNA1 remains bound to microtubules after dilution. (A) Example tubulin dilution profile. The intensity of the background tubulin signal was measured over time before and after tubulin dilution for the control (light) and SSNA1 condition (dark). The time of tubulin dilution is indicated by the arrow. (B) Representative kymograph showing that SSNA1 remains bound to the microtubule following tubulin dilution/washout. Microtubules were first grown with 9 µM tubulin and 5 µM 488-SSNA1 for 60 min, then the reaction solution was washed out and the microtubules were imaged for a further 30 minutes. (C) Control to show that exchanging the reaction solution does not trigger catastrophe in and of itself. Microtubules were grown with 9 µM tubulin for 30 min; the buffer was then exchanged for 9 µM tubulin. The white line on the kymograph indicates the time of tubulin exchange. Note that catastrophe at the plus end does not occur immediately upon exchange and the minus end did not catastrophe over the kymograph duration.

**Figure S6.**
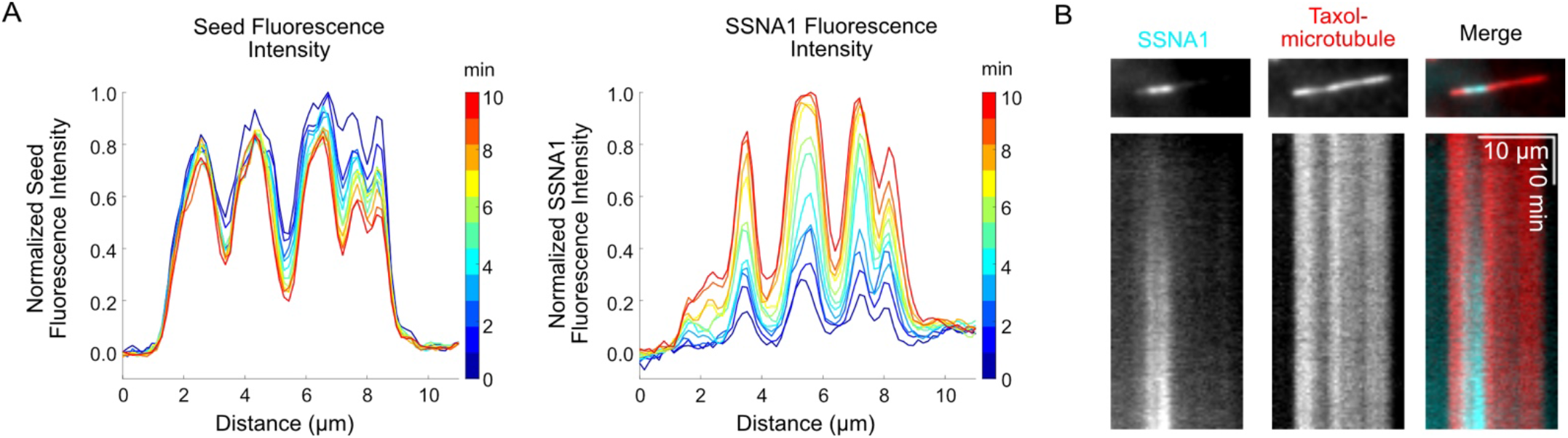
SSNA1 recognizes sites of microtubule lattice damage. (A) Linescan analysis of microtubule seed (left) and SSNA1 (right) fluorescence intensity at 1-minute intervals, color-coded according to time. (B) Taxol-stabilized microtubules were incubated with 5µM 488-SSNA1 and imaged for 60 minutes. Sum projection images of a microtubule (top) and corresponding kymographs (bottom) showing that 488-SSNA1 also recognizes regions of dimmer tubulin fluorescence intensity on the Taxol-stabilized microtubule lattice.

### VIDEO CAPTIONS

**Video S1**. Dynamic microtubule extensions (magenta) growing from GMPCPP-stabilized microtubule seeds (blue) with 10 µM tubulin. Time is in min:s.

**Video S2**. Dynamic microtubule extensions (magenta) growing from GMPCPP-stabilized microtubule seeds (blue) with 10 µM tubulin and 5 µM SSNA1. Time is in min:s.

**Video S3**. Dynamic microtubule extensions (magenta) growing from GMPCPP-stabilized microtubule seeds (blue) with 9 µM tubulin and 5 µM 488-SSNA1. Time is in min:s.

**Video S4**. Dynamic microtubule extensions (magenta) were initially grown from GMPCPP-stabilized microtubule seeds (blue) with 15 µM tubulin alone. At t = 2 minutes, the reaction was exchanged to 15 µM tubulin and 2.5 µM 488-SSNA1. Time is in min:s.

**Video S5**. Dynamic microtubule extensions (magenta) were initially grown from GMPCPP-stabilized microtubule seeds (blue) with 9 µM tubulin alone. At t = 30 seconds, the reaction was exchanged to assay buffer. Time is in min:s.

**Video S6**. Dynamic microtubule extensions (magenta) were initially grown from GMPCPP-stabilized microtubule seeds (blue) with 9 µM tubulin and 5 µM SSNA1. At t = 30 seconds, the reaction was exchanged to assay buffer. Time is in min:s.

**Video S7**. GMPCPP-stabilized microtubule seeds were pre-incubated with SSNA1 storage buffer (control, left) or 1 µM 488-SSNA1 (+ SSNA1, right), and then 200 nM spastin was introduced at t = 0 min. Seeds are in red, SSNA1 is in cyan. Time is in min:s.

**Video S8**. GMPCPP-stabilized microtubules were pre-incubated with 100 nM spastin and then 5 µM 647-SSNA1 was introduced at t = 0 min. Seeds are in red, SSNA1 is in cyan. Time is in min:s.

